# Signaling metabolites spatially organize a multispecies mutualism in the soil microbiome

**DOI:** 10.64898/2026.07.10.737802

**Authors:** Jake A. Drewes, Finlay Warsop Thomas, Julie Bethany, Emily A. Higgins Keppler, Corey Nelson, Suzanne M. Kosina, Trent R. Northen, Heather D. Bean, Ferran Garcia-Pichel

## Abstract

A plant-independent avenue for N_2_-fixation takes place in desert topsoils through a “C-for-N” mutualism between heterodiazotrophs and the cyanobacterium *Microcoleus vaginatus*. These partners come together within a diverse soil microbiome under conditions of N-limitation for the phototroph and C limitation for the heterotrophs. We hypothesized that extracellular chemical signaling might enable partner selection and collocation, though infomolecules shaping inter-microbial architecture were unknown. We show that the complex chemical composition of *M. vaginatus*’ exometabolome depends on its N-limitation status, thus potentially offering information to mutualists. In chemotactic assays, the exometabolome effectively repelled most native soil bacteria, particularly intensely when under N-limitation. Bacterial assemblages circumventing the repulsion were enriched in species that are rare in the soil microbiome, and that functionally resemble mutualistic cyanospheres (showing high N_2_-fixation potential, secretion of urea, and copiotrophy), setting the stage for a working symbiosis. Further, we could reproduce the enrichment of copiotrophs and nitrogen-fixers using mixtures of N-acetylglutamic acid, N-acetylmethionine, indole-3-acetic acid, and 5′-methylthioadenosine, all preferentially released by *M. vaginatus* under N-limitation. These signaling molecules did not result in an enrichment of urea producers, however. The results demonstrate that trans-species communication through specific infochemicals, together with already known quorum-sensing-like intraspecific communication in *M. vaginatus*, act as a tool to organize microbiomes spatially and to attain mutualistic partner specificity in an open, crowded background.

## Introduction

A microbe’s exometabolome is comprised of metabolites of intracellular origin found extracellularly due to loss, secretion, or passive release^1^. Many microbial interactions are enabled through exometabolites, either through syntrophy, where metabolites are exchanged because of their nutritional value, or through specific biological effects they may elicit. One can further distinguish exometabolites influencing the growth of species other than the producer (i.e., allelopathic)^2,3^, such as antibiotics, from those carrying information (signaling molecules, or infochemicals), such as quorum sensing (QS) autoinducers^4^.

In crowded, complex microbiomes, both inter- and intra-species signaling likely mediate emergent system dynamics. Biofilm organization and defense mechanisms, for example, are mediated by intra-specific QS signaling^5,6^. In some cases, QS systems are subject to third-party interference, endowing them with a trans-species dimension^7-9^. Likewise, exometabolite-based trans-kingdom molecular communication between plant or animal host and microbiome has been characterized^10,11^. In the rhizosphere, for example, plant exometabolites chemotactically recruit symbionts to bolster a more advantageous microbiome^12-15^.

Because non-hosted microbiomes also feature interspecies interactions such as division of labor and syntrophy^16-18^, interspecies communication presumably plays as large a part in these dynamics as in hosted microbiomes^19-21^. But its usage among free-living bacteria has yet to be demonstrated, save for the case of QS interference^7^, perhaps because of a lack of appropriate models.

Biological soil crust (biocrust) microbiomes provide such a model. They are photosynthetic microbial communities developing on soil surfaces where plant growth is restricted, and cover some 12% of the terrestrial area^22^. Cyanobacteria often serve as the primary producer^23^, where the filamentous *Microcoleus vaginatus* is the most common^24^, playing a pioneer role in biocrust formation. It, however, cannot fix nitrogen^25^, relying on its proximal heterotrophic microbiome, termed the “cyanosphere”, for its N needs in exchange for fixed carbon^26^. This C-for-N mutualism is crucial for its survival, as its habitat in arid soils is largely N-depleted^27^. The mutualism has been successfully re-created using representative isolates in co-culture^28^. In nature, cyanospheres are variable in specific composition, recruited from the existing bulk soil^29^. They are better defined functionally, as are some rhizosphere microbiomes^30^: intensely enriched in copiotrophic bacteria^29^, possessing a high nitrogen fixation potential (NFP)^29^, and its members secreting urea ^31^. This mutualism therefore offers the opportunity to study a trans-specific interaction in both laboratory and field settings. In fact, exometabolite signaling does play a role through a QS-like system active in N-starved *M. vaginatus* that uses GABA and Glutamate as signals, modifying its own motility behavior^32^. Cyanosphere members interfere with this signaling by producing GABA and Glutamate themselves, when C starved. Yet, it is not known if cyanosphere members display reciprocal motility responses towards *M. vaginatus* or even if the cyanobacterium’s exometabolome itself is affected in an informational way (i.e. involving infomolecules) by N-starvation. In this study, we set out to determine the effects of increasingly intense N limitation on the *M. vaginatus* exometabolome, assess its ability to recruit and assemble a functionally adequate cyanosphere from the bulk microbiome, and identify any infomolecules directly responsible for trans-species communication. We aimed to connect changes in the *M. vaginatus* exometabolome to mechanisms through which it may architect a microbiome optimized for reciprocal nutrient exchange.

## Methods

### Culture Conditions, Media, and Field Sampling

Stocks of *Microcoleus vaginatus* PCC9802 were maintained in BG11 medium^33^ at 23°C with 18– 20 μE m^−2^ s^−1^ of white light illumination under a 14 h/10 h light/dark cycle. For metabolomics, 50 mL cultures were incubated in BG11 modified by varying nitrate concentration from 0 mM to the standard 17.65 mM NO3^-^. Higher concentrations were also tested, but these did not yield informational results and are thus not reported explicitly, although all data are included in Tables S1-2. Below 17.65 mM NaNO3^-^, ionic strength was matched by adding NaCl as needed. Ten independent cultures were grown per concentration. After two weeks of incubation, spent medium was filter-sterilized (0.22 µm pore size; Pall Corporation) and aliquots taken (40 mL for soluble compounds and 10 for volatiles) and stored at -80°C until analysis. As a source of microbiomes, natural biocrusts were collected from Jornada Basin soils (New Mexico, USA) using Petri plates to capture the top 1 cm^34^, air dried, and stored in darkness until experimentation. The dominance of *M. vaginatus* was confirmed microscopically^35^.

### Soluble metabolomics

Thirty mL of spent media were lyophilized and then resuspended in 1 mL mass spectroscopy grade methanol, vortexed twice for 10 s, bath sonicated in ice water for 20 min, and then centrifuged at 10,000 rcf, for 5 min at 10°C. Supernatants were dried by vacuum concentration (Thermo Savant SpeedVac SPD111V) and stored at -80°C until analysis. Just prior to LCMS analysis, samples were resuspended in 150 µL ice-cold methanol containing internal standards. Vortexing, sonication, and centrifugation were repeated and supernatants were filtered through 0.22 µm PVDF microcentrifugal filter tubes at 10,000 rcf for 10 min at 10°C. Filtrates were transferred to amber glass vials and capped. Metabolites were separated using hydrophilic interaction liquid chromatography and analyzed using a Thermo Scientific hybrid quadrupole-orbitrap Q Exactive mass spectrometer (instrument parameters are also available in **Table S1**). Targeted annotations were performed using Metatlas^36-38^. Identifications were verified according to the Metabolomics Standards Initiative (MSI) minimum reporting standards^39^. Untargeted features from negative mode were extracted using MZmine 2^40^. Analytes were retained for further untargeted analysis (**Table S1**) only if the maximal value across all samples was 10,000 units greater than the maximal value across controls.

### Volatilomics

Volatile organic compounds (VOC) were sampled using solid-phase microextraction (SPME)^41^ from 2 mL from each spent medium dispensed into 10 mL gas chromatography (GC) headspace vials with septum magnetic screw caps. A 1 cm, 23 Gauge Divinylbenzene/Carboxen/Polydimethylsiloxane fiber was placed in the headspace and equilibrated with the sample. Fibers were then retrieved and metabolites released by heat within the chromatography setup. VOCs were analyzed by two-dimensional GC coupled with time-of-flight mass spectrometry (GC×GC–TOFMS). Parameters used for desorption, chromatography, mass spectrometry, data processing, and peak alignment (performed using the Leco ChromaTOF^®^ software with the Statistical Compare package; Leco Corp.) are in **Table S4**. Prior to data analysis, chromatographic contaminants and artefacts were removed from the peak table. For VOCs that were detected in at least 60% of replicates within a sample group, missing values were imputed using the replicate Random Forest algorithm from the R MetabImpute package (0.1.0)^42^. VOCs failing that 60% threshold had all replicates imputed to 0. The relative abundance of compounds was normalized using probabilistic quotient normalization (PQN)^43^. Analytes were retained for eventual analysis based on the following criteria: (i) an intraclass correlation coefficient (ICC) greater than or equal to 0.4, (ii) significantly greater in abundance in any sample type than the blank medium control using the Wilcoxon rank-sum test with Benjamini-Hochberg correction and an alpha of 0.05, and (iii) arithmetic mean of any biological replicate peak areas at least two-fold greater in abundance than the blank medium control. A base two logarithmic transformation was then applied for downstream processing.

### Microbiome Chemotaxis Assay

Chemotaxis of biocrust microbiome members towards *M. vaginatus* exometabolomes, compared to defined, artificial control solutions, was assayed using pipette tips filled with solution and brought into contact directly with the top 2 mm of biocrust soil for a duration of 48 h (**Fig. 1**), after which the solutions were retrieved and analyzed. One hour prior to the start of the assay, biocrusts were soaked with N-free BG110 ^33^ medium to soil saturation to reactivate quiescent bacteria. The entire assay set-up was then enclosed in a partially sealed plastic bag to help maintain soil saturation, with periodic re-wetting with medium as needed. For soluble metabolomes, tips contained 300 µL volumes. For volatiles, an air gap of approximately 100 µL was placed between 100 µL of test solution and 100 µL of control artificial medium, which acted as a “receiving solution” in contact with the soil (**Fig. 1**). This ensured that chemotactic effects were caused by compounds that could equilibrate through the air gap. NaCl at 15% (w/v) was added to the test solutions just prior to loading to enhance volatility of its contents^44,45^. A description of effectively sampled biocrust volumes and other potential confounding factors is in **Supplementary Methods**.

**Figure 1:**
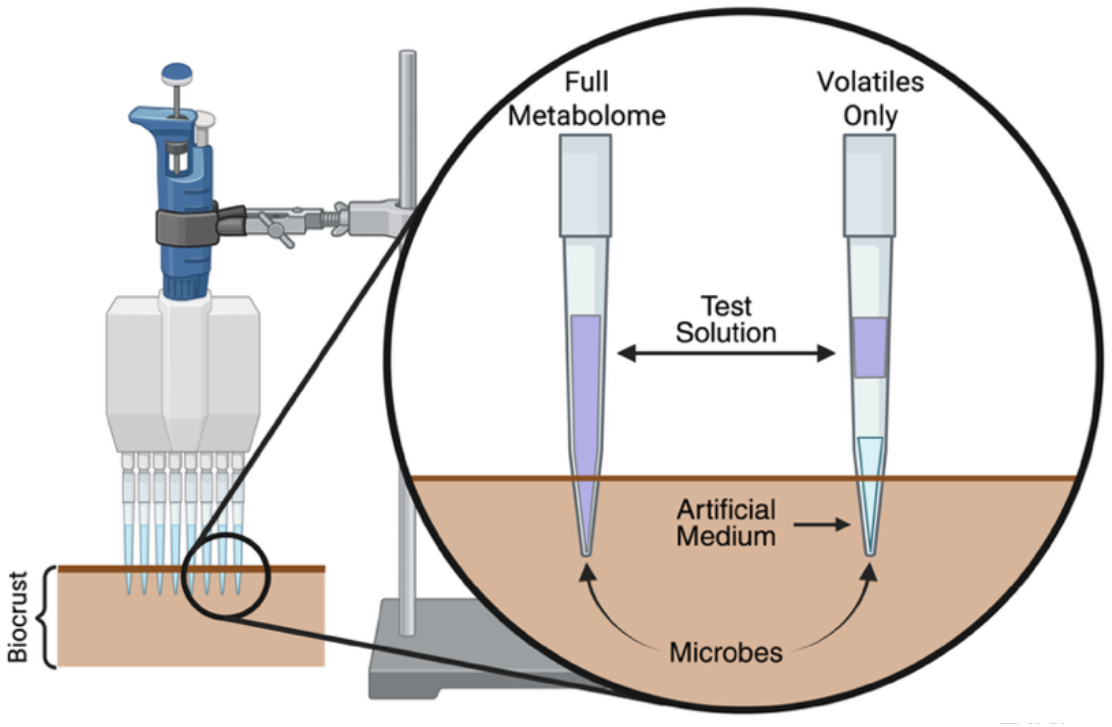
Chemotaxis assay set-up. A stand held a multichannel pipette in place so that micropipette tips would sit inside a saturated biocrust sample. The tips were filled with 300 µL of test solution (for soluble exometabolome or soluble infochemicals ; left tip in circle) or fwith 100 µL of solution, a 100 µL and 100 µL of standard control medium, the latter remaining in contact with the crust (right tip in circle).

Test solutions included filter-sterilized spent medium from N-starved *M. vaginatu*s, and the spent medium of unstarved cultures. Chemically-defined control solutions consisted of BG11_0,_ amended with tryptic soy broth and glucose to reach 0.25 g C L^-1^ and 0.04 g N L^-1^ (pH = 6.02). The spent medium of N-starved cultures contained 0.03 g C L ^-1^ and 0.16 g N L^-1^ (pH = 8.43), whereas the unstarved contained 0.10 g L C ^-1^ and 0.22 g L N ^-1^ (pH = 8.43) as measured on a Shimadzu TOC-L analyzer. The control provides a common reference to compare across experiments. For controls, 21 independent tip assays on 12 occasions with different biocrust samples were conducted and combined prior to analysis to yield a total of 15 determinations. Based on normality and interquartile range analyses, two of these determinations were deemed to be outliers and were excluded from downstream comparisons. To test the effect of specific compounds or their combinations, we spiked the control solution with up to 100 µM. This kept pH shifts < 0.7 units.

### Determination of Microbiome Size, Composition, and NFP

Community DNA in the assay solutions and soil samples was extracted using the Powersoil Pro extraction kit (QIAGEN, Hilden, Germany) according to manufacturer’s instructions. DNA extracts were PCR-amplified, targeting the V4 region of the 16S rRNA gene using 515F/806R primers^46^, and amplicons sequenced on a MiSeq sequencer (Illumina). Sequences were demultiplexed using the DADA2 plugin of QIIME2^46,47^. After trimming for quality control and discarding singletons, taxonomic classification was initially performed using SILVA^48^.

The number of bacteria and their microbial N_2_-fixing potential (NFP) in the attracted communities were assessed through quantitative real-time PCR (qPCR) as 16S rRNA and *nifH* gene copies per mL of test solution. The ratio of *nifH* to 16S rRNA counts was taken as a measure of NFP. The primer set 338 F (5′-ACTCCTACGG GAGGCAGCAG -3′) and 518 R (5′-GTATTACCG CGGCTGCTGG -3′)^49^ was used for 16S rRNA genes, and the PolF (5′-TGCGAYCCSA ARGCBGACTC-3′) and PolR (5′-ATSGCCATCA TYTCRCCGGA-3′)^50^ set for *nifH*. Reactions were carried out in triplicate using PerfeCTa SYBR Green FastMix (Quantabio, Beverly, MA, USA) in an ABI ViiA 7 thermocycler (Applied Biosystems, Foster City, CA, USA) with previously published protocols^26^. For statistical comparisons, the ratio of *nifH* to 16S rRNA gene copies was log-transformed to account for variance in homogeneity and normality.

### Community diversity and composition

16S rRNA sequences from a previous study of the same locality and biocrust type^28^, gathered using identical DNA extraction and sequencing parameters, were used as a reference for the composition of cyanosphere and bulk soil microbiomes. Chao and Shannon alpha diversity indices were assessed using the R^51^ package phyloseq^52^. Bray-Curtis distances of sample groups were produced using QIIME2 and principal coordinate analyses (PCoA) were plotted using the qiime2^53^ R package. Sequence identity for the ASVs representing the top 70% of total reads per treatment group was corroborated using NCBI BLAST^54^. ASVs were categorized as oligotrophs or copiotrophs according to phylogenetic placement.

### Urea Production

Urea concentrations in the chemotaxis assay solutions were determined immediately following biocrust incubation with the Amplite® Colorimetric Urea Quantitation Kit (AAT Bioquest) using a UV-2600i Plus spectrophotometer (Shimadzu) and normalized to the number of copies of the 16S rRNA gene. The resulting ratios were log-transformed prior to statistical testing.

### Statistical analyses and data presentation

The similarity in exometabolome profiles was explored using principal component analysis (PCA) and hierarchical clustering analysis (HCA), with samples as observations and absolute peak intensities (which were first mean-centered and scaled to unit variance) as variables using R version 4.2.1^51^. 3D PCA and PCoA plots for exometabolomics and beta diversity data were made with the R package plotly^55^ (4.10.1). HCA heatmap was made with the R package pheatmap^56^ (1.0.13). Treatment effects in chemotaxis assays, urea production assay, alpha diversity, and copiotrophy analyses were assessed through ANOVA, with Tukey’s HSD post-hoc tests if data were normally distributed (Shapiro-Wilk test) and had equal variance (Levene’s test). If either failed, the Kruskal-Wallis test was used, with Conover-Iman post-hoc testing. Plots were created using ggplot2^57^ (3.5.0). All test results are in **Table S3**.

### Data availability

Soluble metabolomic raw data files for positive and negative polarities and processed untargeted negative data are available at the Global Natural Products Social Networking 2 (GNPS2)^58^ database (https://gnps2.org/status?task=61b19eae694743ed9a56295ff18ff4cb).

Raw volatile metabolomic data are available at the NIH Common Fund’s National Metabolomics Data Repository (NMDR) website, and at the, metabolomicsworkbench.org^59^, with project ID PR003006 and study ID ST004738 (DOI: 10.21228/M8427G).

Raw 16S rRNA sequencing data are available at NCBI under Bioproject PRJNA1459394.

## Results

### Nitrogen starvation alters the composition of the *M. vaginatus* exometabolome

We analyzed both the soluble and volatile compounds found in spent media of *M. vaginatus* grown at different nitrate concentrations (**Fig. 2**). For the soluble exometabolome, an untargeted analysis using HILIC-ESI-MS/MS detected 2,436 features among all samples (**Table S1**). Using an atlas of known mz, retention time, and MS2 values, we could identify 84 metabolites (**Fig. S1**). 82% of which were nitrogenous, including amino acids and derivatives, and nucleosides/nucleobases. Other commonly identified compounds were carboxylic acids and carbohydrates. For the volatilome, an untargeted analysis using GC×GC–TOFMS of solid-phase microextracts yielded 85 features among all samples (**Table S2**).

**Figure 2:**
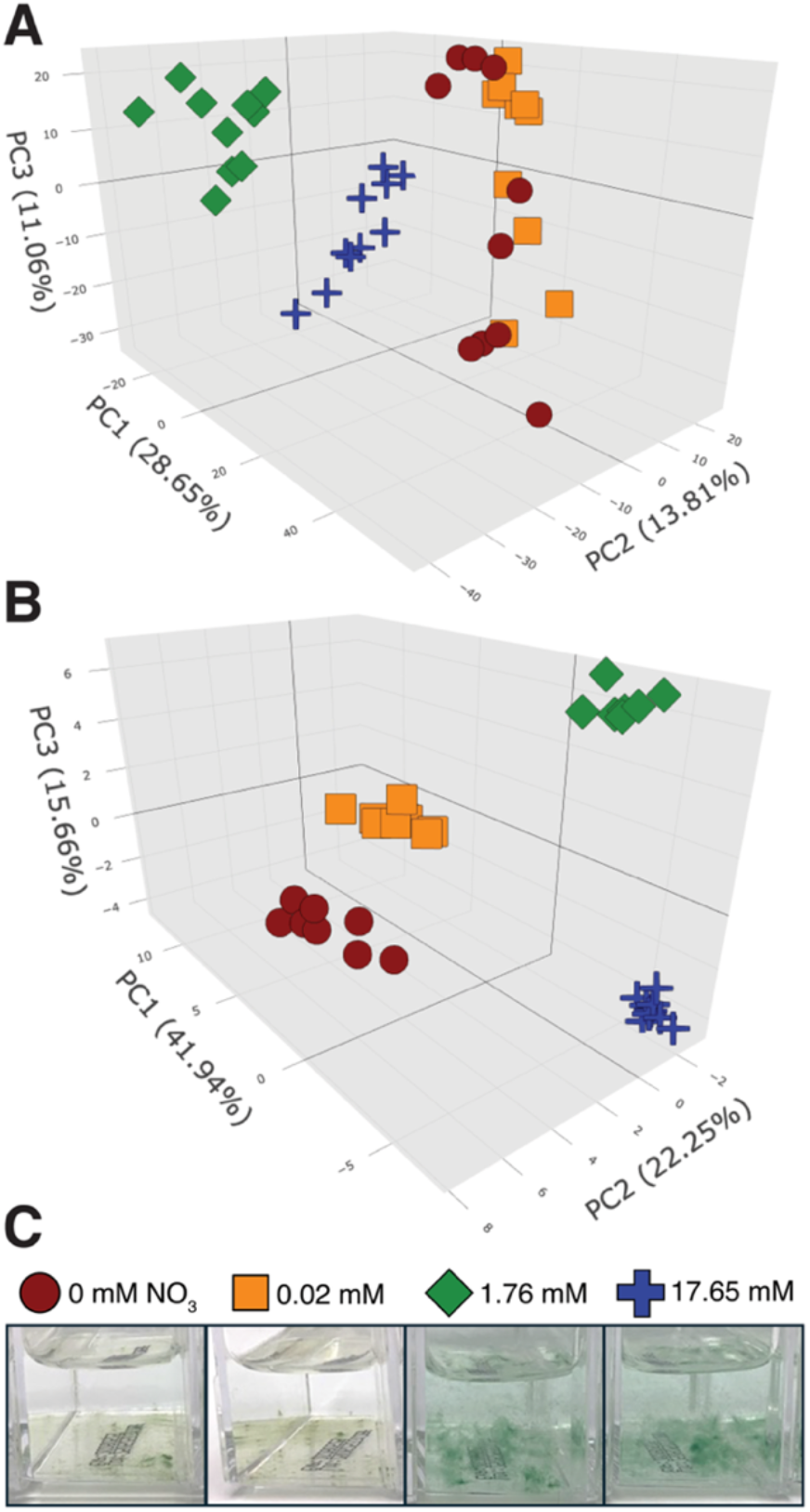
Principal component analyses (PCA) of the soluble **(A**; n=10) and volatile **(B**; n=7-9) *M. vaginatus* exometabolome fractions when grown under defined nitrogen concentrations, with their respective phenotypes **(C)**.For PCAs, biological replicate cultures were used as observations and either 2,436 soluble features **(A)** or 82 volatile features **(B)** were used as variables. For soluble features, the first three principal components encompass 54% of the variance and for volatiles 80%.

Principal component analyses (PCA) of the untargeted datasets revealed differences in composition with N status (**Figs. 2A-B**). In both exometabolomes, replicates were self-similar and the two lowest N-availability treatments (0 and 0.02 mM NO_3_^-^) clustered together and on the opposite side of PC1=0 from the rest (**Figs. 2A-B**). This mirrored cellular symptomology, where low N cultures showed the chlorosis typical of strong N starvation, unlike those in higher concentrations (**Fig. 2C**). This demonstrates that the exometabolome of *M. vaginatus* cells carry potential information about its nutrient status in the form of a differential chemical composition, both in its soluble and volatile fractions.

Potential N-starvation signaling molecules within the exometabolome can be predicted to increase in concentration as N decreases. A search for specific compounds among the targeted soluble dataset revealed only a few conforming to this pattern (**Fig. 3**): one tetrasaccharide, one trisaccharide (neither one more precisely identified), one nucleobase derivative (5′-methylthioadenosine) and several amino acid derivatives (pipecolic acid, N-acetyl glutamic acid, N-acetyl methionine, hydroxyphenyllactic acid, indoleacetic acid). Of these, N-acetylglutamic acid (NAG), indole-3-acetic acid (IAA), N-acetylmethionine (NAM), and 5′-methylthioadenosine (5′-MTA) were available commercially and were used for further study.

**Figure 3:**
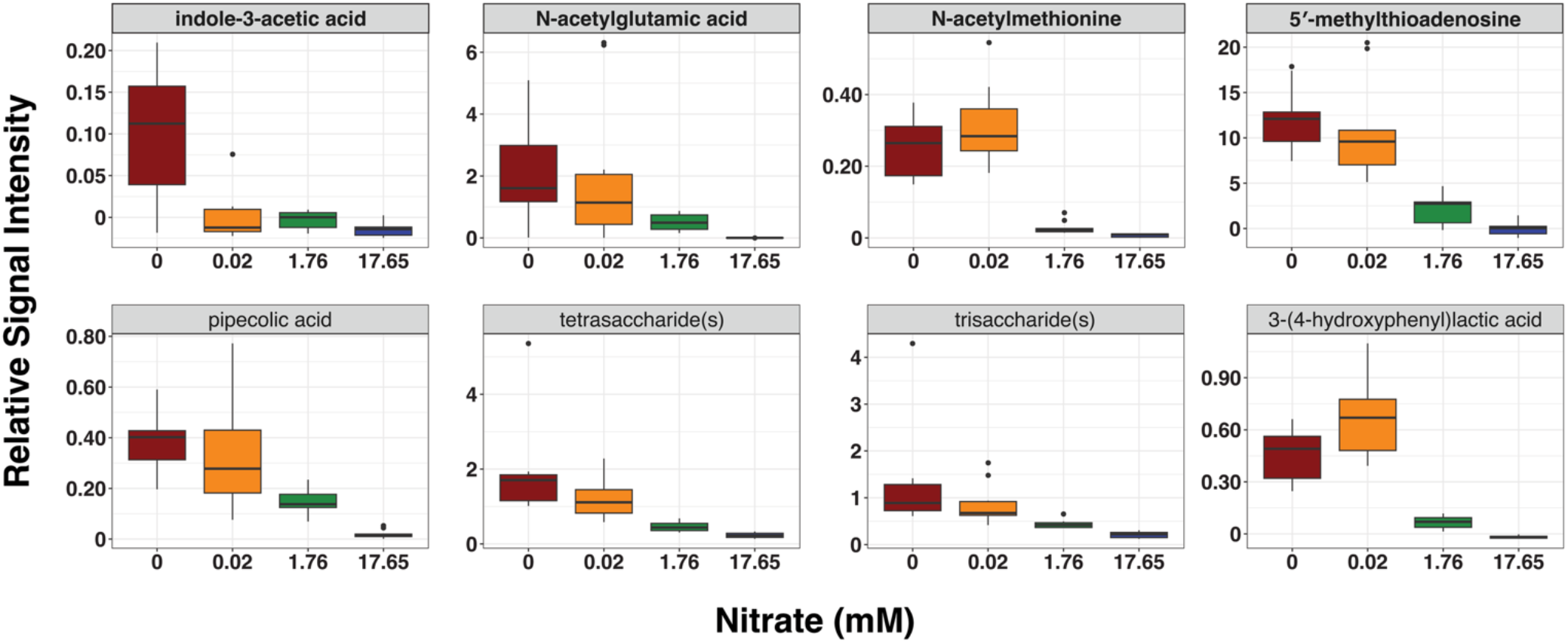
Relative signal intensity of potential infochemicals detected in the soluble exometabolome at different nitrate concentrations. For each compound, the signal intensity at the two lowest and two highest N concentrations was significantly different (n=10; see **Table S3** for significance values and test details). The four compounds used for further study are labelled in bold type.

Thus, the exometabolome of *M. vaginatus* is complex, depends on its nutrient status with respect to N, and contains specific metabolites potentially useful as tell-tale signs of N-starvation.

### Chemotactic effects of the *M. vaginatus* exometabolome on the biocrust microbiome

Chemoattraction assays (**Fig. 1)** of natural biocrust microbiomes towards *M. vaginatus* exometabolomes showed significant effects against a standardized control solution. The exometabolome of starved cultures attracted much fewer bacteria than the control solution as judged by the concentration of 16S rRNA gene copies (a proxy for bacterial numbers), by two orders of magnitude (**Fig. 4; top left)** (p<0.0001). Average 16S rRNA gene concentrations in the controls reached 10^8^-10^9^ mL^-1^, which is similar to those existing in typical biocrust microbiomes^23^, whereas only 10^6^-10^7^ reached the starved exometabolome solution. Thus, the control solutions essentially equilibrated in numbers with the source microbiome, while the exometabolome prevented such equilibration. Exometabolomes from unstarved *M. vaginatus* cultures had a similar effect, albeit less intense (and only marginally significant; p=0.09). The control solution was a defined standard to compare between treatments and experiments. It had a higher C content (0.25 g C L^-1^) than the starved (0.03 ± 0.002 g C L^-1^ ; n=6) or unstarved (0.07 ± 0.02 g C L^-1^; n=6) exometabolomes, which may help explain why more bacteria were attracted. However, the magnitude of the effect in bacterial numbers seems incommensurate with the smaller difference in C content. Alternatively, the contrasting results may be explained by net exometabolome chemorepellence, which we address in the next section.

**Figure 4:**
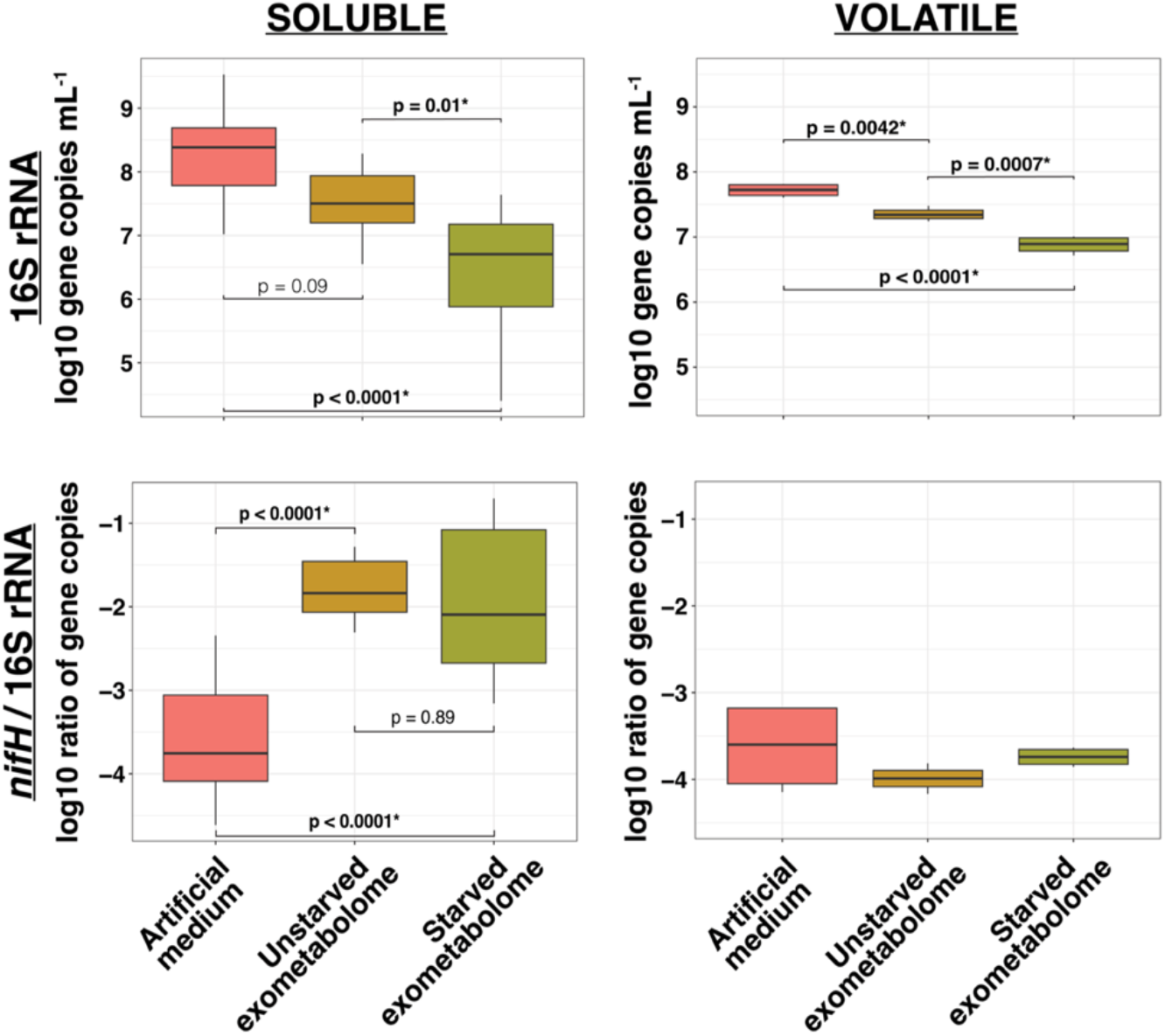
16S rRNA concentrations and N_2_-fixing potential (NFP) in the solutions used in the chemotaxis assay. The effects of the full *M. vaginatus* exometabolome and the subset of volatile components of the exometabolome were assessed with relation to nitrogen starvation and compared to an artificial medium control. For the volatilome, n=8 for each treatment. Treatments for the full soluble exometabolome experiment varied: artificial medium n=17, unstarved n=8, starved n=13.

With respect to the NFP of communities attracted (assessed as paired *nifH* to 16S rRNA copy ratios), both exometabolomes were significantly more enriched in diazotrophs than the control (p<10^-5^), (**Fig. 4; bottom left**). This cannot be a consequence of a differential N content, as the control (0.04 g N L^-1^) was N-poorer than the starved (0.17 ± 0.02 g N L^-1^; n=6) or unstarved (0.20± 0.02 g N L^-1^; n=6) exometabolomes and should have attracted more N_2_-fixers. For comparison, the typical (log) NFP of bulk biocrust microbiomes is between -3.5 and -5, depending on origin, whereas that of *M. vaginatus* cyanospheres hovers between -2.2 and -2.5^29,60^. Hence, the NFP of attracted communities were within the cyanosphere range or slightly higher, whereas those of controls did not differ from bulk biocrusts. We could not demonstrate statistically, as we may have predicted, a difference in NFP enrichment between the starved and unstarved exometabolomes. Increases in the NFP of exometabolomes resulted from a combination of 16S rRNA and *nifH* tallies, but the former was most influential in the outcome, as *nifH* copies did not always increase significantly (**Fig. S2A**).We conclude that *M. vaginatus’s* soluble exometabolome depresses the bacterial numbers attracted over controls, simultaneously allowing an enrichment in diazotrophs to cyanosphere-like levels in the resulting communities.

We also tested if the volatile fraction of the exometabolome could elicit similar effects. The concentration of bacteria attracted to the control were within the range obtained for controls in **Fig. 4** (i.e., equilibrated with the source soil). Because replicate variability here was much smaller, we could detect significant treatment decreases in 16S rRNA with respect to controls, even when they were smaller (less than an order of magnitude) than those seen with the soluble exometabolome (**Fig. 4; top right**). Unlike with the soluble exometabolome, here we tested the superimposition of the volatilome over a background of control solution **(Fig. 1)**. Therefore, the decrease in bacterial populations exerted by the volatilomes must have been due to a chemorepellent effect, which was much stronger with the volatilome of starved cells. Hence, as with the full exometabolomes, the volatile fractions also exerted a net repulsive effect on total bacterial numbers that was stronger under N-starvation, though overall weaker than that of full exometabolomes. In volatilome tests, *nifH* gene concentrations (**Fig. S2B**; p=0.018) were in fact significantly lower than in control solutions. The effects on bacterial numbers and potential diazotroph numbers largely cancelled each other so that the NFP of populations attracted to volatilomes did not differ significantly from controls (p=0.3), and none of the treatments resulted in an NFP higher than those typical of bulk biocrusts (**Fig. 4; bottom right**). We conclude that the volatilome contributes a net repulsive effect on soil bacteria, though much less intensely than the full exometabolome, but does not contribute significantly to altering the resulting NFP. Specific tests that follow, therefore, focus on the soluble exometabolome.

### Tracing chemotactic effects to compounds

We then conducted chemotaxis assays towards solutions of the control medium amended with varying concentrations of four compounds characteristic of the starved exometabolome: NAG, IAA, NAM, and 5′-MTA. The concentrations represent initial conditions within the tips, likely decreasing in time and with distance from the tip by diffusion. While 5′-MTA had no effects, the others were clearly chemotactically active, though not in the direction we could have predicted. NAG, IAA, and NAM at 100 µM all increased the number of bacteria attracted, some significantly (**Fig. 5A**). Spiking control medium with 100 µM NAG, IAA, or NAM resulted in significant NFP decreases over controls, contrary to expectations, with no significant effects for 5′-MTA.

**Figure 5:**
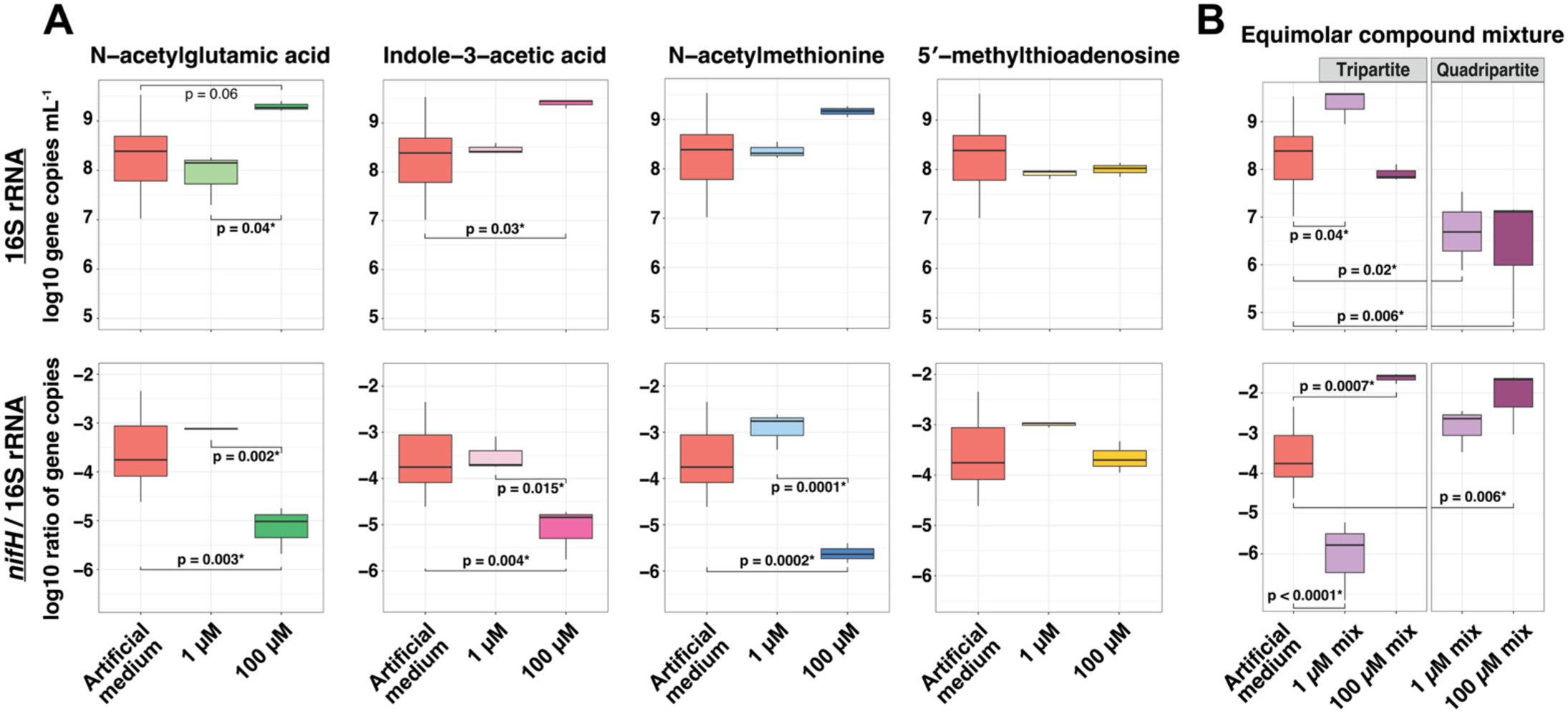
16S rRNA concentrations and N_2_-fixing potential (NFP) of communities attracted to the artificial medium (n=17) versus (**A)** artificial media supplemented with one of four putative infochemicals (NAG, IAA, NAM, 5′-MTA) added at increasing concentrations, or (**B**) artificial medium supplemented with equimolar mixtures of either a quadripartite mixture of the four putative infochemicals NAG, IAA, NAM, 5′-MTA) or a tripartite mixture of the three compounds that individually elicited an crease in NFP (NAG, IAA, NAM). (all n=6)

To investigate synergistic effects, we also tested the compounds in an equimolar quadripartite mixture (NAG, IAA, NAM, 5′-MTA) and a tripartite mixture that excluded 5′-MTA, for which we had seen no bioactive effects as a single compound. We found strong synergisms in a concentration-dependent manner (**Fig. 5B**). The tripartite mixture at 1 µM (0.33 µM each) attracted more bacteria than controls but had no effects at 100 µM. The quadripartite mixture had a strong net-repelling effect, by two orders of magnitude, regardless of concentration (**Fig. 5B**). With respect to NFP, the tripartite mixture resulted in lowered NFP at 1 µM, and higher at 100 µM, both effects two orders of magnitude different than controls. The quadripartite mixture by contrast, increased the NFP at both concentrations, by about one order of magnitude at 1 µM (though non-significantly with p=0.19 and by two at 100 µM. Synergistic interactions among infochemicals are obviously important and complex. The strong synergistic effects of 5′-MTA seem to involve most prominently the general repulsion of bacteria (i.e. compare the 1 µM differences between the tripartite and quadripartite solutions in **Fig. 5B**, top panel) but play a minor role, if at all, in raising the NFP. For this reason, subsequent tests to characterize communities attracted used only the tripartite solution.

We conclude that these compounds were bioactive alone or in combination, but also that only their combined synergistic activity reproduced the net repulsion of bacteria and the enrichment in NFP seen with the raw exometabolome, reversing the effects determined in isolation. Because the results demonstrate a net chemorepellent effect of these compounds, as they were tested over a background of the control solution, they also speak for chemorepellence as a logical explanation for the results of soluble exometabolomes in **Fig. 4**.

### The *M. vaginatus* exometabolome attracts communities functionally similar to cyanospheres

Since the vehicle for mutualistic nitrogen transfer by heterodiazotrophs to *M. vaginatus* is urea^31^, the ability to secrete urea constitutes a stringent test for the presence in an assemblage of potential mutualists. Urea secretion is otherwise quite rare among bacteria, especially diazotrophs. We therefore determined the levels of urea found in the communities attracted to tests and control solutions at the end of the assay. Both the starved and unstarved full exometabolome communities displayed a ten-fold higher ratio of urea produced per 16S rRNA gene copy than the controls or the control solution amended with a tripartite mixture of NAG, IAA, and NAM (**Fig. 6A**), although the starved exometabolome did not do this more intensely than the unstarved exometabolome. This demonstrates the selective effect of the exometabolome on recruitment of potentially mutualistic bacteria but also suggests that compounds other than those used in the mixture must be behind this specific functional enrichment.

**Figure 6:**
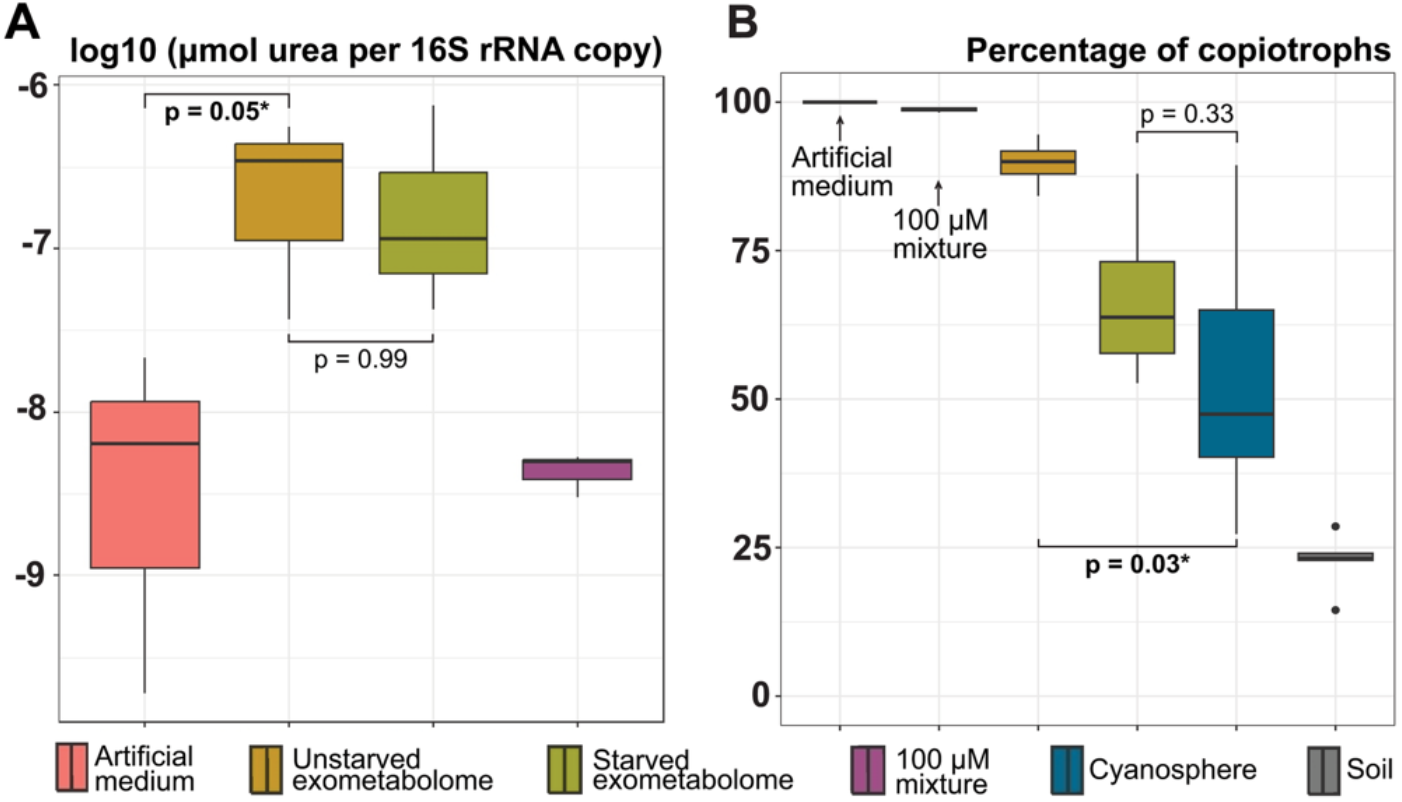
**(A)** Production of urea by the communities attracted to test solutions, normalized to the number of 16S rRNA copies in them. (**B**) Proportion of copiotrophs in the top 70% most abundant ASVs in the communities. (all n=6)

A categorization of the Amplicon Sequence Variants representing the top 70% of reads in attracted communities based on 16S rRNA gene phylogeny (**Fig. 6B**, detailed in **Table S5**) showed that artificial control medium and the control medium spiked with the tripartite compound mixture were either entirely or almost entirely made up of copiotrophs. Typical cyanospheres, while variable, show a less marked copiotrophic composition. Copiotrophy in communities attracted to exometabolomes was non-significantly (starved, p=0.33) or significantly (unstarved, p=0.03) more prevalent than that of natural cyanospheres. These effects are most parsimoniously explained not by differences in chemical composition, but by differences in organic carbon content, as the percentage of copiotrophs mirrors the concentration of organics in the respective test (and control) solutions. Attracting a cyanosphere enriched in copiotrophs may simply be the result of a higher concentration of organics close to *M. vaginatus* against a lower background in the bulk soil solution, and not necessarily the result of infochemical effects. However, the assembled communities with the closest percentage of copiotrophs to native cyanospheres were those of the N-starved exometabolome.

### Community assembly is affected by exometabolites

The previous analyses told us about effects on bacterial numbers and community function, but not on bacterial species composition or diversity. To assess this, we analyzed community composition analyses through 16S rRNA gene sequencing. This showed that the bacterial assemblages attracted to test solutions were diverse, with average (Shannon, S) diversity in the range of 3-5 (**Fig. 7A**). For comparison, the reference cyanospheres averaged 3.8, whereas the bulk soil they originated from hovered around 4.9. Only the artificial control medium assembled significantly less diverse communities than that of the soil, while the others did not differ. Bacterial richness (Chao) was much more variable than S diversity. The highest richness by far was attained in assemblages attracted to the starved exometabolome (642), much higher than those attracted to the unstarved counterpart (360). The starved exometabolome was richer than the soil itself, but the unstarved was not. As with S, the control solution attained the lowest bacterial richness (111). The 100 µM tripartite mixture resulted in an assemblage of intermediate richness between those in the exometabolome tests. That differences are made more apparent with Chao than with S speaks for the importance of rare species in responding to differential attraction. For example, N-starvation resulted in attraction of almost double the number of ASVs over the unstarved version, but only insignificant increases in S, implying that starvation leads to the attraction of many more “rare” bacterial species; so rare, in fact, that they raise the Chao index over that of bulk soil. Thus, many of the ASVs attracted must not have been detectable in the bulk soil analyses. We deduce that the general repulsion of bacterial numbers effected by the exometabolome did not concurrently decrease bacterial diversity, but in fact increased it by recruiting ASVs from the biocrust “rare biosphere”.

**Figure 7:**
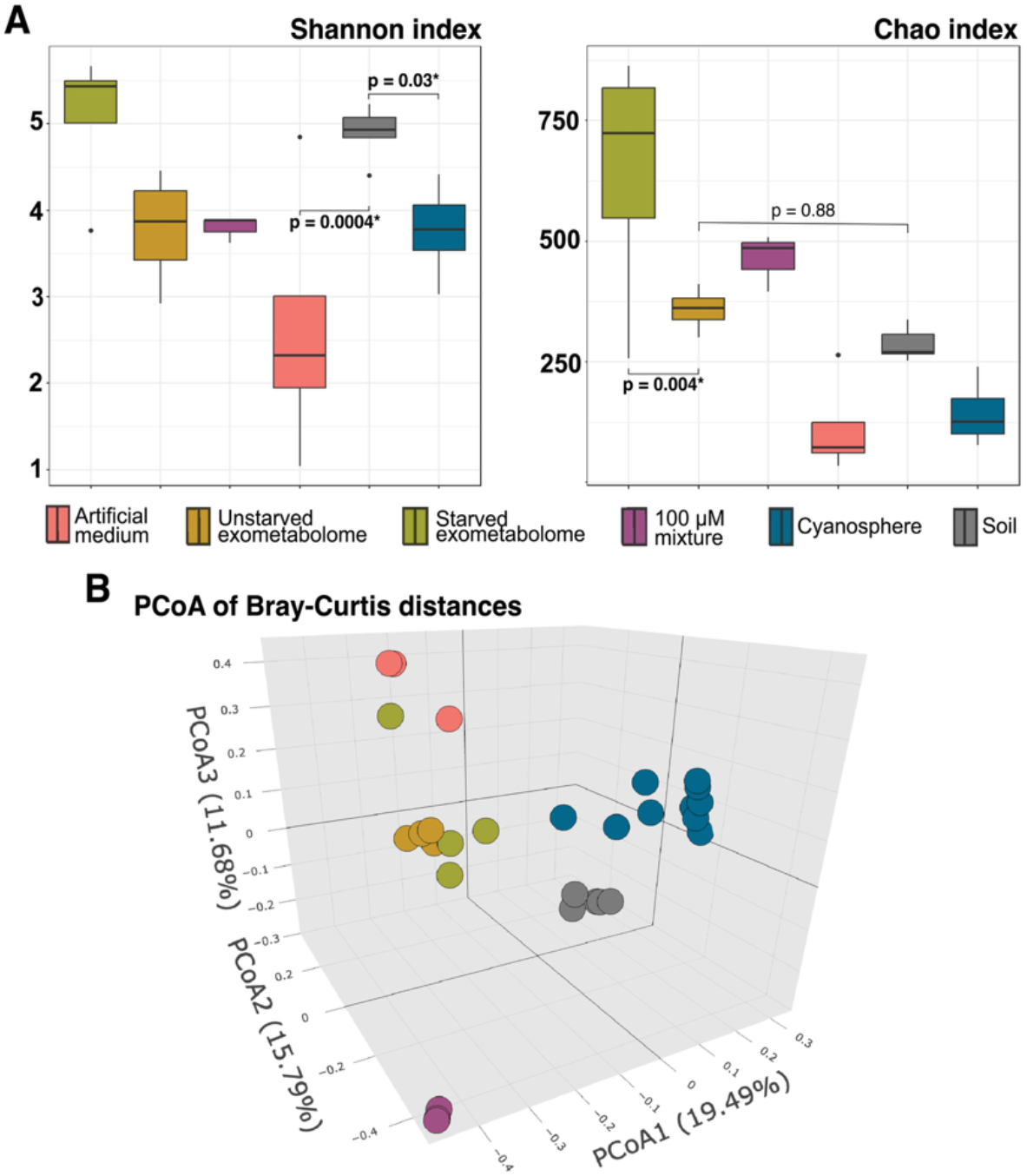
Diversity analyses of communities attracted to test solution compared to soil and cyanosphere communities. (**A**) Alpha diversity analysis using Shannon and Chao indices. (**B**) Beta diversity analysis depicted with principal coordinate analysis of Bray-Curtis distances. Each point represents a biological replicate.

The communities attracted to the test solutions had unique compositions (**Fig. 7B**), though a compositional differentiation between starved and unstarved exometabolomes was not patent. In all cases communities assembled in the assays differed from those of the biocrust soil and mature cyanosphere communities as well. That species composition of exometabolome-enriched communities failed to resemble those of native cyanospheres implies that the process of cyanosphere assembly is not complete at the stage of initial recruitment.

## Discussion

The use of infomolecules in interspecies microbial interactions, especially those involving microbiome organization, is poorly understood. We provide phenomenological evidence from biocrust bacteria that cross-species extracellular signaling is a driver of chemotactic responses enabling the formation of a keystone mutualism. We showed that the exometabolome composition of *M. vaginatus*, the cyanobacterial partner, reflects the availability of N, particularly when under N limitation of growth. Because N-limitation primes the cyanobacterium for mutualism^26^, its exometabolome under N-limitation bears the potential to act as a signal for its readiness to enter C-for-N mutualism. Consistently, the exometabolome selectively repelled bacteria from native biocrust assemblages regardless of nutrient status and involving both soluble and volatile fractions. However, the intensity of the effect was much stronger under N-limitation. As the chemorepellent effect of the volatilome was weaker than that of the soluble fraction, effects can be traced largely to soluble compounds under the assay conditions used. The activity of volatiles, however, may become particularly relevant in extending the range of action, in that they can have a much longer diffusional distance than soluble metabolites in undersaturated soils, as has been shown in rhizosphere communities^61^, but neither long-range attraction nor unsaturated soils were tested here.

The selective exometabolome repulsion of biocrust bacteria also enriched for communities of increased copiotrophic nature, high NFP, and urea-production capacity, which are precisely the functional traits that characterize native cyanosphere communities. The enrichment in copiotrophs can be explained in part by a non-specific chemoattraction to solutions rich in organic compounds, as it also occurred in control solutions, and not necessarily on the bioactivity of specific infochemicals. Even then, this is not inconsequential, as this is likely to play out in nature as well. The relative enrichment in nitrogen fixers cannot be explained in the same way, as controls had a lower N content than exometabolomes. Neither can the enrichment in urea producers, which must be mediated by the presence of bioactive compounds. Hence, chemotactic effects, general or specific, resulted in communities that resemble in these respects the natural mutualistic cyanosphere, supporting our initial hypothesis.

Four compounds produced almost exclusively under N-limitation (NAG, IAA, NAM and 5′-MTA), exerted a net repulsion of bacteria similar to that seen with whole exometabolomes, but only synergistically. Their effects were neutral or net-attractive when tested in isolation. While their role as infochemicals in generalized chemoattraction or repellence is clear, the net effect depends on each other’s presence. A combination of these metabolites can thus be considered a means for cyanosphere recruitment. Because the unstarved exometabolome, which had only traces of these infomolecules, retained some bioactivity, the four signaling compounds identified cannot constitute the totality of trans-species chemotactic signals. Others surely remain to be identified. Further, that the three compounds that increased NFP did not lead to an increase in the urea secretion capacity of the bacterial communities recruited (**Fig. 6A**) indicates that other exometabolites must exert this particular effect. Candidates must be sought in N-starvation independent compounds, in that the enrichment was elicited by starved and unstarved exometabolomes alike.

Two of these infochemicals have been previously implicated in plant-microbe symbioses. NAG produced by root nodule bacteria induces root hair wall growth and stimulates cortical cell division in clover^62^, as needed for the nitrogen-fixing symbiosis. Soybean plants inoculated with IAA knock-out rhizobacteria also produce fewer nodules^63^. As parallels between the cyanosphere and rhizosphere continue to come to light^64^, it is noteworthy that these infomolecules support nitrogen-fixing mutualisms in both. 5′-MTA, largely implicated in synergistic bacterial repellence here, is known to disrupt QS systems through the inhibition of 5′-methylthioadenosine/*S*-adenosyl homocysteine nucleosidases (MTANs), enzymes responsible for the biosynthesis of some autoinducers^65,66^.

The experimental approaches used here may have been suboptimal in some respects, leading to a loss of sensitivity in our assays. First, by necessity, only motile bacteria will be able to respond to signals, when demonstrably some cyanosphere bacteria are non-motile (i.e. *Pseudarthrobacter* spp.)^26^. Second, the patchy nature of natural microbiomes surely contributed to large replicate variability, with an ensuing loss of sensitivity. Given that patchiness in biocrust microbial composition manifests itself strongly at cm scales^60^, precisely the sampling scale in the assays (around 1 cm^3^, see **Supplementary Methods**), we implemented high replication (>8), but in some cases it may have been insufficient. In-tip growth likely contributed to the final population tallies in the assay tips, though this is unlikely to have introduced bias (see **Supplementary Methods**). Finally, in counting *nifH* copies (and calculating the NFP) in a community, particular phylotypes may have been missed because the corresponding PCR primers are not universal^67^.

The results obtained in this work, together with recent reports of QS-like systems in *M. vaginatus*, and QS-interference by several of its heterotrophic mutualists^32^, contribute to a parsimonious mechanistic explanation for the fine-scale spatial organization of biocrust microbiomes (**Fig. S4**). Our model starts with a random distribution of microorganisms in the soil, and the onset of N limitation. This elicits the release of GABA/Glu in *M. vaginatus*, increasing the frequency of motility reversals, which causes an aggregation of multifilament bundles. Initially, bundles will aggregate randomly along the EPS sheath, where steep diffusional gradients of informational exometabolites (and photosynthate) will form, repelling most bacteria in their surroundings, and shifting the character of bundle-proximal populations to one of increased copiotrophy, NFP, and urea secretion capacity. This enriched local population stands to benefit most directly from photosynthate exudation, providing in turn a local source of urea. Upon this optimized collocation, functionally capable mutualists can be further selected by syntrophic compatibility. For as long as these mutualist populations receive growth-limiting amounts of photosynthate, they will also produce GABA/Glu, further enhancing recruitment of cyanobacterial filaments into the bundle, and shifting the dynamic location of bundles along the sheath preferentially to where large populations of mutualistic exist. If recruitment of cyanobacteria to the bundle is excessive, levels of GABA/Glu will reach its high-concentration negative feed-back loop, which will elicit bundle disaggregation. The combined intra- and interspecies molecular chatter driving bacterial motility behavior thus seems an important tool in attaining the spatial segregation that shapes the biocrust microbiome into distinct cyanosphere regions, optimized for mutualism, and the remaining bulk.

## Supporting information

Supplemental Table 1: Soluble metabolomics parameters and results

Supplemental Table 2: Volatile metabolomics results

Supplemental Table 3: Statistical Test Results

Supplemental Table 5: Copiotrophy determinations

## Supplemental Materials

**Figure S1**: Hierarchical clustering analysis of the soluble exometabolites produced by *M. vaginatus* grown at different nitrate concentrations

**Figure S2**: *nifH* gene copy concentration in the communities attracted to chemotaxis assays **Figure S3**: Directly determined 16S rRNA gene copy concentration and concentration of urea in communities attracted to chemotaxis assays, as used to calculate specific production of urea **Figure S4**: Cartoon diagram for a working signaling model

**Table S1**: Excel workbook containing LCMS parameters, LCMS targeted annotation of 85 compounds with evidence, and 2,436 features from the untargeted LCMS analysis, detected in liquid *M. vaginatus* cultures

**Table S2**: Excel worksheet of the 89 VOCs detected in the untargeted analysis of the headspace of liquid *M.vaginatus* cultures

**Table S3**: Excel worksheet of statistical test results for all intergroup differences

**Table S4**: Parameters for headspace SPME, GC×GC-TOFMS analysis, and volatile data processing and alignment

**Table S5**: Excel worksheet of the genera of heterotrophs considered copiotrophs or oligotrophs for the trait-based categorization of ASV’s in 16S rRNA communities

## Supplementary Methods

## Acknowledgements

This project was supported in part by Arizona State University School of Life Sciences New and Bold Funding to FGP and HB, as well as the NSF grants DEB 2129537 and DEB 2425143 to FGP. This work was also funded by the Office of Science Early Career Research Program, Office of Biological and Environmental Research, of the U. S. Department of Energy under contract number DE-AC02-05CH11231 to TN.

## Author contributions

FGP and HDB conceived of the study; FGP, HDB, JAD, FWT, JB, and CN designed the experiments; JAD, FWT, JB, and CN collected the data; JAD, FWT, JB, CN, EAHK, SK, and TN processed and analyzed the data; JAD, FWT and FGP wrote the manuscript; all authors edited the manuscript and approved the final version.

## Conflicts of Interest

T.R.N. is a founder of two non-profits, Prosper Soils and Bioaligned Labs with prior approval from LBNL. We declare that the research was conducted in the absence of any commercial or financial relationship that could be construed as a potential conflict of interest.

## Notes

https://gnps2.org/status?task=61b19eae694743ed9a56295ff18ff4cb

https://www.metabolomicsworkbench.org/data/DRCCMetadata.php?Mode=Study&StudyID=ST004738&StudyType=MS&ResultType=5

### Competing Interest Statement

The authors have declared no competing interest.

