## Supplemental Information for "Signaling metabolites spatially organize a multispecies mutualism in the soil microbiome"

**Figure S1**: Hierarchical clustering analysis of the soluble exometabolites produced by *M. vaginatus* grown at different nitrate concentrations. Analysis was done for the forty replicates and 84 soluble exometabolites using Manhattan distance and ward.D linkage for compounds. Relative abundance of each compound is depicted by a color gradient from blue (indicating lower abundance) to red (higher abundance). The eight compounds of interest are boldened (see **Fig. 2**) and the four compounds that were used for further experimentation in this study are highlighted with a red rectangle.

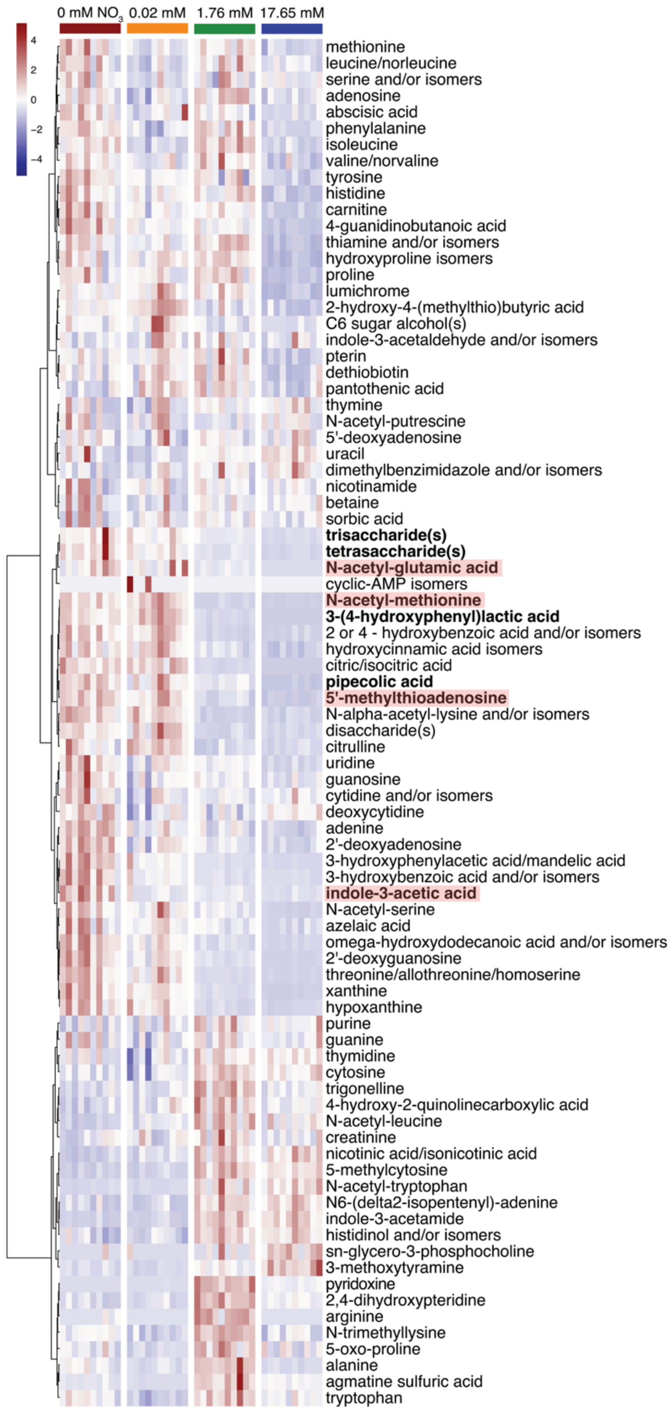

**Figure S2**: *nifH* gene copies concentration in the communities attracted to chemotaxis assays. Assays include artificial medium, spent medium from *M. vaginatus* under either normal conditions or under nitrogen starvation, and artificial medium spiked with possible infochemicals at different concentrations. (A): the full exometabolome as shown in **Fig. 4**; (B): the volatilome only as shown in **Fig. 4**; (C-F): the four possible infochemicals individually as shown in **Fig. 5A**; (G): the compounds in mixture as shown in **Fig. 5B**.

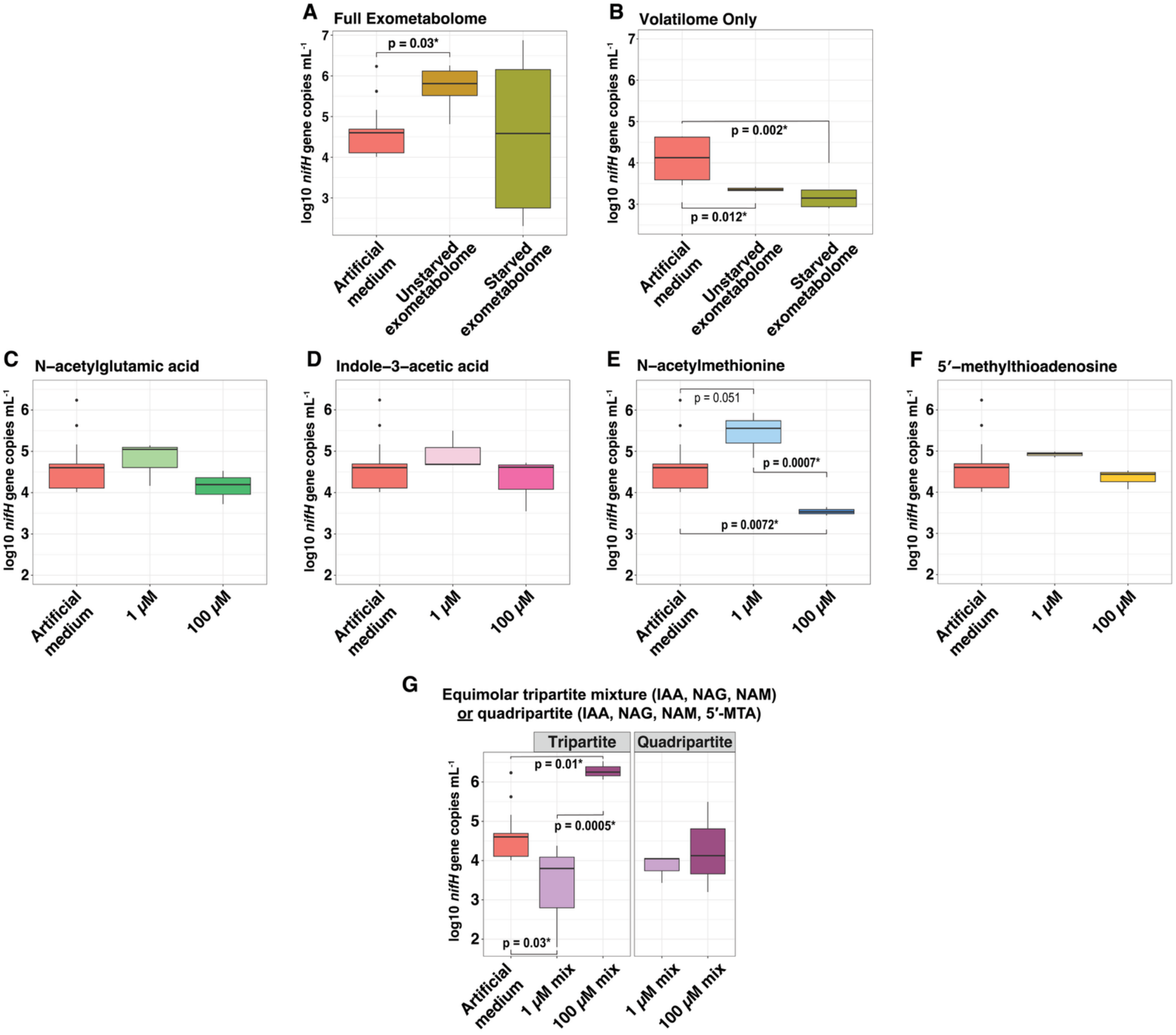

**Figure S3**: Directly determined 16S rRNA gene copy concentration and concentration of urea in communities attracted to chemotaxis assays, as used to calculate specific production of urea in **Fig. 6A**.

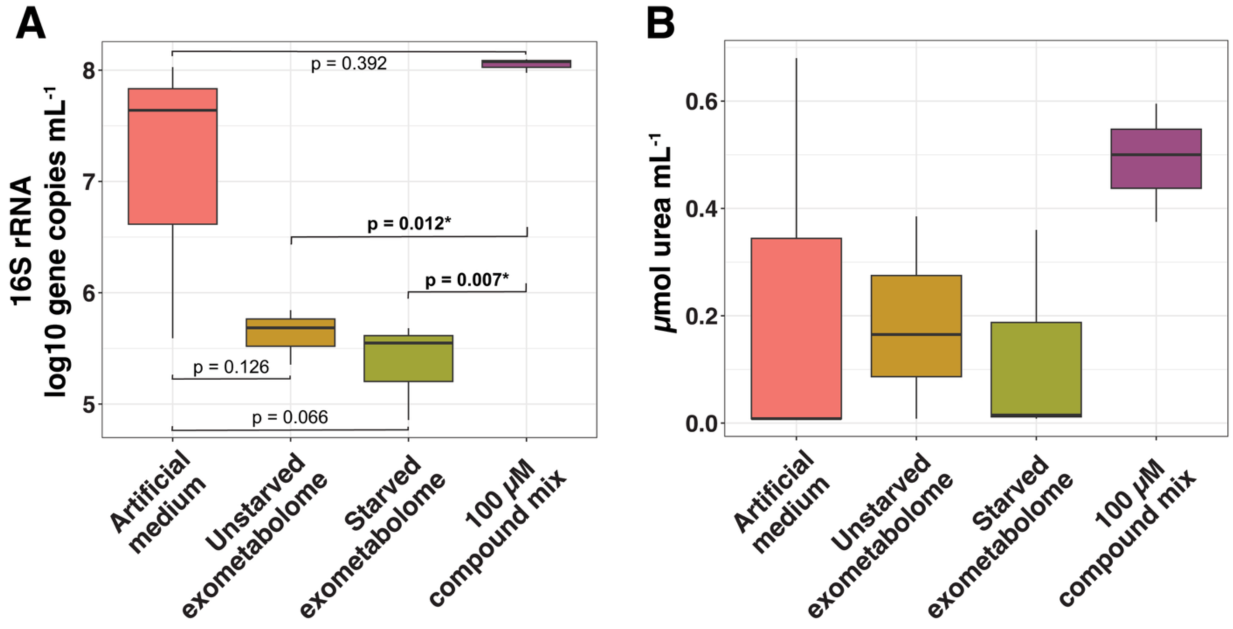

**Figure S4:** Cartoon diagram for a working signaling model. Upon nitrogen limitation and increased bundle formation, *M. vaginatus* secretes distinct soluble and volatile metabolites. At closer proximities to *M. vaginatus*, soluble metabolites, including IAA, NAM, NAG, and 5′-MTA help to shape the surrounding community by enriching for $N_2$-fixers, urea producers, and copiotrophs. Since volatile metabolites are capable of passing into the air phase and moving farther than soluble compounds, these metabolites may act to generally repel all bacteria and minimize possible competition.

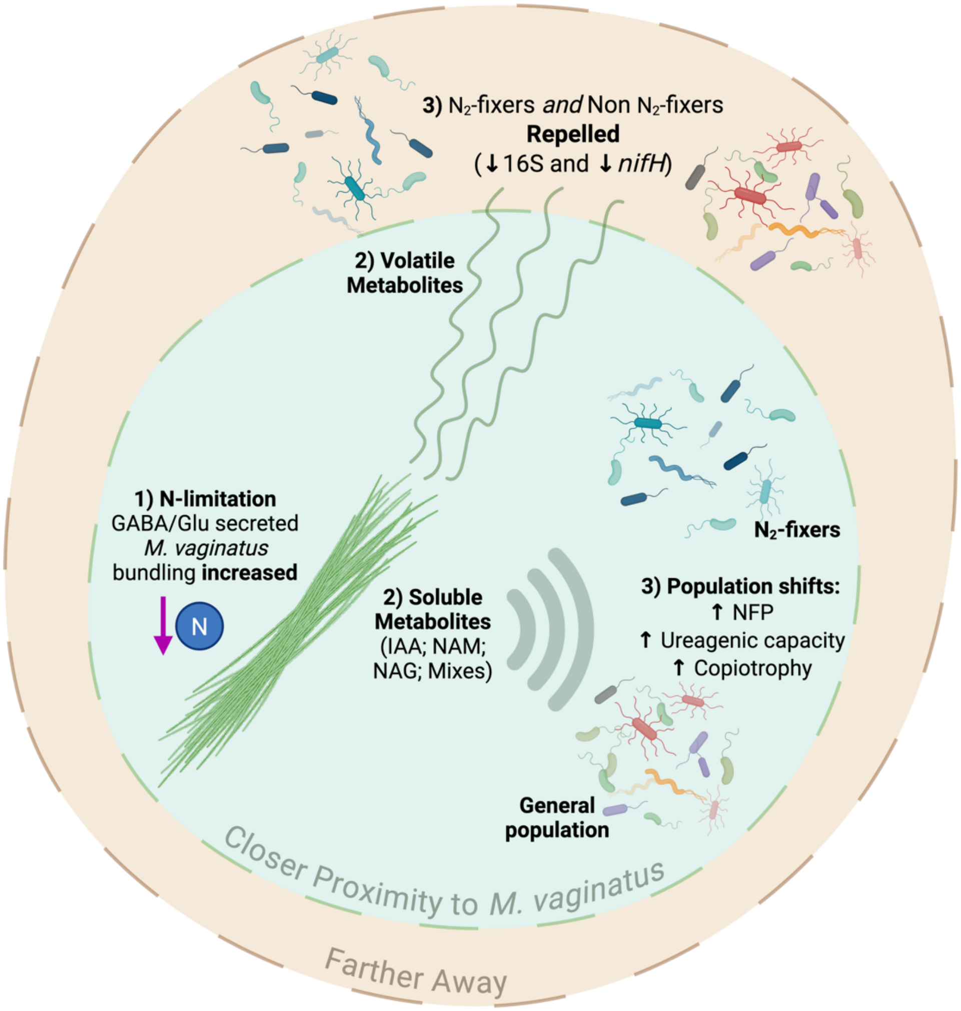

**Table S1 See Supplementary Excel File.** Workbook containing the following pages: 1) LCMS parameters, 2) table of the 85 annotated compounds from targeted LCMS analysis with evidence including ID levels according to the Metabolomics Standards Initiative (MSI) minimum reporting standards[46], and 3) table of the 2,436 features from untargeted LCMS analysis. For each compound the following information is reported: a compound identification number assigned by the data processing software, average compound m/z, and average retention time in seconds. For each of the sample groups and media blank, the number of biological replicates each compound was detected in is reported along with the mean relative peak height and standard deviation. The sample group "Uninoculated BG11o + 52.95 mM $NO_3$-" was included in the experiment by mistake and was not used for any downstream analyses.

**Table S2: See Supplementary Excel File.** Table of the volatile features detected in the headspace of liquid *Microcoleus vaginatus* PCC9802 cultures when cultured in BG11 medium with various amounts of nitrate supplied. For each compound the following information is reported: a volatile organic compound (VOC) identification number assigned by the data processing software, average first- and second-dimension retention times ($^1t_R$ and $^2t_R$, respectively), and the observed first-dimension retention index on 624-Sil stationary phase. For each of the sample groups and media blank, the number of biological replicates each compound was detected in is reported along with the mean relative peak area and standard deviation.

**Table S3: See Supplementary Excel File.** Statistical test results by figure and individual sample group.

**Table S4.** Parameters for headspace SPME, GC×GC–TOFMS analysis, and volatile data processing and alignment

| AUTOSAMPLER METHOD | |
|---|---|
| Instrument description | Gerstel® MPS Pro® |
| Software description | Gerstel® Maestro® (version 1.5.3.2) |
| **Sampling Parameters** | |
| Cooled tray temperature | 4°C |
| Solid-phase microextraction (SPME) | Manufacturer: Supelco®<br>Fiber type: PDMS/CAR/DVB (2 cm) |
| Incubation time | 5 min |
| Agitator parameters | Temperature: 50°C<br>On time: 10 s<br>Off time: 1 s<br>Speed: 600 RPM |
| Vial penetration | 21 mm |
| Extraction time | 10 min |
| Injection penetration | 67 mm |
| Desorption time | 180 s |
| **Thermal Desorption Parameters** | |
| Initial temperature | 50 ºC |
| Ramp rate | 720 $ºC·min^{-1}$ |
| End temperature | 220 ºC |
| Hold time (min) | 3.00 |
| Transfer temperature | 240 ºC |
| Transfer temperature mode | Fixed |
| Desorption mode | Splitless |
| Sample mode | Retain Tube – Standby Cooling |
| Standby temperature | 50 ºC |
| **Inlet (CIS) Parameters** | |
| Initial temperature | -80 ºC (cryo cooling) |
| Equilibrium time | 0.20 min |
| Initial time | 0.0 min |
| Ramp rate | 12 $°C·s^{-1}$ |
| End temperature | 250 °C |
| Hold time | 3.00 min |
| Heater mode | Standard |
| Liner | Glass liner, non-baffled w/glass wool |
| **TWO-DIMENSIONAL GAS CHROMATOGRAPHY METHOD** | |
| Instrument description | Agilent® 7890B |
| Software description | Leco ChromaTOF (version 4.72.0) |
| Column configuration | Column 1: Rxi®-624Sil MS, 60 m × 0.25 mm × 1.4 $\mu$m<br>Column 2: Stabilwax®, 1 m × 0.25 mm × 0.5$\mu$m |

| | |
|---|---|
| Carrier gas | Helium, 2 mL·min$^{-1}$ (constant) |
| Front inlet type | Gerstel® |
| Front inlet mode | Splitless |
| Front inlet septum purge flow | 1 mL·min$^{-1}$ |
| Front inlet septum purge time | 300 s |
| Front inlet purge flow | 50 mL·min$^{-1}$ |
| Front inlet total purge flow | 52 mL·min$^{-1}$ |
| Oven equilibration time | 5 s |
| Primary oven temperature ramp | Initial temperature: 35 °C<br>Initial time: 0.5 min<br>Ramp rate: 5 C·min$^{-1}$<br>Final temperature: 230 °C<br>Hold time: 5 min |
| Secondary oven temperature offset | +5 °C (relative to primary oven) |
| Modulator temperature offset | +15 °C (relative to secondary oven) |
| Modulation timing | Modulation period: 2.00 s<br>Hot pulse time: 0.50 s<br>Cold pulse time: 0.50 s |
| Transfer line temperature | 250 °C |
| **MASS SPECTROMETRY METHOD** | |
| Instrument description | LECO® Pegasus® 4D |
| Use GC method total time for MS method total time | Yes |
| Acquisition delay | 180 s |
| Filament active time | 180 s to end of run |
| Start mass/End mass | 35/400 |
| Acquisition rate | 100 spectra·s$^{-1}$ |
| Optimized voltage offset | +50 V |
| Electron energy | -70 eV |
| Ion source temperature | 250 °C |
| **DATA PROCESSING METHOD** | |
| Software description | LECO® ChromaTOF® with Statistical Compare (version 4.71.0.0) |
| Baseline tracking/Offset | Entire run/0.5 (through middle of noise) |
| Data points averaged for smoothing | Auto |
| First dimension peak width | 8 slices |
| Mass spectral match required to combine | 600 |
| Second dimension peak width | 0.1 |
| Min. subpeak signal-to-noise (S/N) for | 6 |
| Integration approach | Traditional |
| Peak finding | S/N: 50<br>Number of apexing masses: 2 |
| Mass spec libraries for searching | NIST 2011 |
| Library identity search mode | Normal |

| | |
|---|---|
| Library search mode | Forward |
| Minimum molecular weight allowed | 35 |
| Maximum molecular weight allowed | 550 |
| Mass threshold (0-998) | 10 |
| Mass to use for area/height calculation | Unique mass |
| Alignment analyte match criteria | Spectral match mass threshold: 10<br>Minimum spectral similarity match: 600<br>Max. number of modulation periods apart: 3<br>Max. retention time difference (s): 0.2<br>S/N for second peak find: 5 |
| Criteria for inclusion of analytes | Min. number of samples that contain analyte: 1<br>Min. % of samples in class that contain analyte: 50 |

**Table S5: See Supplementary Excel File.** Table of the genera of heterotrophs considered copiotrophs or oligotrophs for the trait-based categorization of ASV's in 16S rRNA communities.

**Supplementary Methods**

In the configuration used for the chemotaxis assays, the effectively sampled soil volume can be estimated from the average diffusional distance of signal molecules, $x$, from the tip opening. For small molecules with the diffusion coefficient D of around $5x10^{-10}$ $m^2$ $s^{-1}$ and a 48 h incubation time, t, the radial diffusional distance, given by $x^2$=2Dt, comes to 1.3 cm at the end of the incubations. The corresponding time-averaged radius is 0.65 cm, which corresponds to that of a sampled volume sphere of 1.1 $cm^3$.

The distance potentially covered by motile migrating bacteria with typical swimming speeds in 48 h is well above 100-200 cm. This implies that bacterial immigration rates within a sampled volume with maximal distances of 1.2 cm, should not have constrained the size of the immigrating populations.

In-tip growth may have contributed to the final numbers of bacteria reached in the assays. To assess the contribution of growth vs immigration to the final tallies, we modeled exponential growth as a function of immigration rate, *I*, doubling time, *d*, and the time of residence in the tip since arrival and parametrize this model with literature data. The yield and growth rates of natural mixed bacterial communities in liquid media provided with external C is determined by the concentration provided: final yields are a direct function of concentration by approximately 5 x $10^5$ cells $ml^{-1}$ per mM C provided, and doubling times are independent of concentration (doubling times of 17-23 h) above 1 mM C (0.01 g C $L^{-1}$) provided[1]. In our case then, with test solution around 0.01-0.25 g C $L^{-1}$, growth rates should approximate maximal values, and biomass yields should not have been achieved at 48 h of incubation but have remained around 1-2 x $10^5$ cells $ml^{-1}$. With a content of 10 copies of the 16S rRNA copies per cell[2] of modal bacteria with a volume of 0.9 µm$^3$ [3] , only our lowest experimental tallies approximated this cell concentration, so it is unlikely that growth was curtailed by an exhaustion of substrate during the assays. Setting *d* = 20 h, a copy of the 16S rRNA gene reaching the tip at the beginning of the assay will have contributed 5.6 copies after 48 h. One arriving at 8 h into the assay (with 40 h to grow) will only contribute 4, and one arriving at 28 h will contribute 2. For a constant *I*, the number of cells contributed by *in situ* growth follows an exponential decay with time of arrival, where the time integral of that function through the assay period provides the total number from all arrivals. For example, if *I* =1 copy h$^{-1}$ and *d* = 20 h, then the total contribution is 126.6 copies, of which 48 immigrated and 78.6 grew in-tip. In this case, the percentage of final counts attributable to growth is 45%. Because the total contribution depends directly and linearly on *I* (as 126.6 x *I)* but so does the total immigrated bacteria (as 48x *I*), the percentage is independent of migration rates, and the % of counts attributable to growth is only dependent on *d*. With *d* ranging from 17 to 23 h$^{-1}$*,* the percentage of in-tip grown copies in our tallies should be between 52 and 39%. Thus, any treatment tallies that significantly differ by less than two-fold could have been unduly influenced by growth rather than immigration, but, in hindsight, this was never the case.

Differential growth rates among immigrant bacteria, however, may have also contributed to an apparent differential enrichment (i.e., to species composition of the communities attracted), increasing the relevance of copiotrophs.


1 Eiler, A., Langenheder, S., Bertilsson, S. & Tranvik, L. J. Heterotrophic bacterial growth efficiency and community structure at different natural organic carbon concentrations. *Appl Environ Microbiol* **69**, 3701-3709 (2003). https://doi.org/10.1128/AEM.69.7.3701-3709.2003

2 Gonzalez-de-Salceda, L. & Garcia-Pichel, F. The allometry of cellular DNA and ribosomal gene content among microbes and its use for the assessment of microbiome community structure. *Microbiome* **9**, 173 (2021). https://doi.org/10.1186/s40168-021-01111-z

3 Secaira-Morocho, H., Chede, A., Gonzalez-de-Salceda, L., Garcia-Pichel, F. & Zhu, Q. An evolutionary optimum amid moderate heritability in prokaryotic cell size. *Cell Rep* **43**, 114268 (2024). https://doi.org/10.1016/j.celrep.2024.114268
